# Paradoxical effects of Zim3, a CRISPRi effector, on human iPSC-cardiomyocyte electrophysiology

**DOI:** 10.1101/2023.06.10.544468

**Authors:** Julie L. Han, Yuli W. Heinson, Maria R. Pozo, Weizhen Li, Emilia Entcheva

## Abstract

We show that Zim3, when used as Zim3-KRAB-dCas9 effector in interference CRISPR, without any guide RNAs, paradoxically upregulates key cardiac ion channel genes in human induced pluripotent stem-cell-derived cardiomyocytes, iPSC-CMs, responsible for healthy resting membrane potential, repolarization of the action potential and electrical transmission of signals. These were found to yield expected functional enhancements consistent with a more mature iPSC-CM phenotype, with potentially desirable properties.

## Introduction

Zinc Finger imprinted 3 (Zim3 or ZNF657/264) is a protein-coding gene in humans, found at low levels in testis, brain, skeletal muscle and skin(1). As other ZNF-KRAB (Krueppel associated box) domains, which bind to DNA and through complexing with KAP1 (KRAB-associated protein 1) affect chromatin organization(2), Zim3-KRAB may be involved in the regulation of transcription at multiple genomic loci. In the human heart, the expression of Zim3/ZNF657/ZNF264 has not been examined in detail.

## Results

We analyzed RNAseq data in human adult heart (left ventricle, LV) from the Gene Tissue Expression (GTEx) database(3) and in human iPSC-CMs, **Figure 1**. The Zim3 gene identifier was only expressed in a very small fraction of the samples, at negligible levels, both in the LV and in our iPSC-CMs. ZNF657 was not expressed in either group of samples. The ZNF264 gene was most consistently expressed – in all adult heart samples and in all iPSC-CM samples, but also at very low levels. For comparison, we also show the gene expression for KCNH2, encoding for the main repolarizing K+ channel in cardiomyocytes. To quantitatively compare the two datasets, we normalized the transcripts per million (TPM) by the respective geometric mean, GM, across the transcriptome in each sample and displayed log10 of the ratios(4). The adult human heart (LV) has a higher ZNF264 expression than the human iPSC-CMs, however both have very low expression overall.

**Figure 1.**
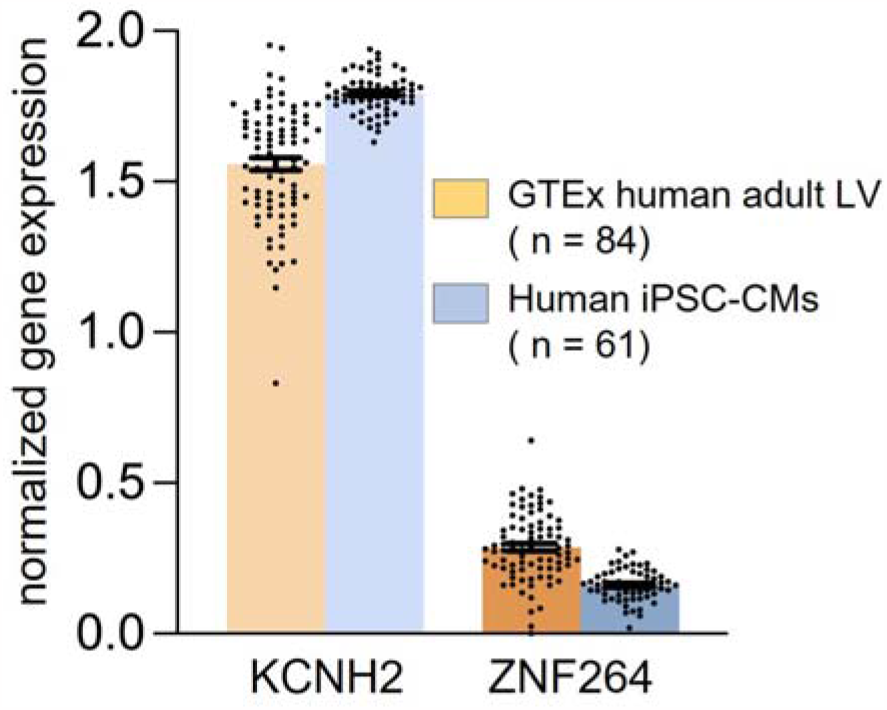
Endogenous gene expression of Zim3/ZNF657/ZNF264 in human adult heart (left ventricle, LV) from GTEx and in human iPSC-CMs, along a well-expressed cardiac gene, KCNH2. The Zim3 and ZNF657 gene identifiers were either not expressed or expressed in only a small subset of samples at negligible levels. For ZNF264 and for KCNH2, we show log10 of normalized TPM, log10(TPM/GM).

Recently, Zim3-KRAB has been identified as a potent effector (repressor) in combination with dCas9 for interference CRISPR (CRISPRi) gene regulation studies(5). Indeed, using Zim3-KRAB-dCas9 adenoviral delivery in combination with suitable gRNAs in post-differentiated induced pluripotent stem cell-derived cardiomyocytes (iPSC-CMs), we have shown robust suppression of the expression of KCNH2 with concomitant functional consequence(6). While performing control experiments, we observed unexpected effects of simply expressing Zim3-KRAB-dCas9 alone on cardiac electrophysiology, without the addition of any gRNAs or with the use of scrambled/control gRNA. Namely, we found Zim3-KRAB-dCas9 expression to significantly upregulate the mRNA expression of key cardiac ion channels: KCNJ2, encoding for the Kir2.1 protein – inward rectifier K+ channel, contributing to the maintenance of healthy negative resting membrane potential; KCNH2, encoding for the rapid delayed rectifier that controls action potential duration, APD; and GJA1, encoding for Cx43 – the main gap-junctional protein in ventricular cardiomyocytes; for Cx43 this was also confirmed at the protein level. Using all-optical electrophysiology, **Figure 2a**, we corroborated physiologically relevant functional effects, **Figure 2c-e**, expected for upregulated KCNH2 (APD shortening in paced conditions), **Figure 2c, f**; upregulated KCNJ2 (significantly reduced spontaneous firing), **Figure 2d, f**; and upregulated GJA1 (conduction velocity increase), **Figure 2e-g**. In combination, these electrophysiological changes are perceived as desirable and a signature of more mature iPSC-CMs.

**Figure 2.**
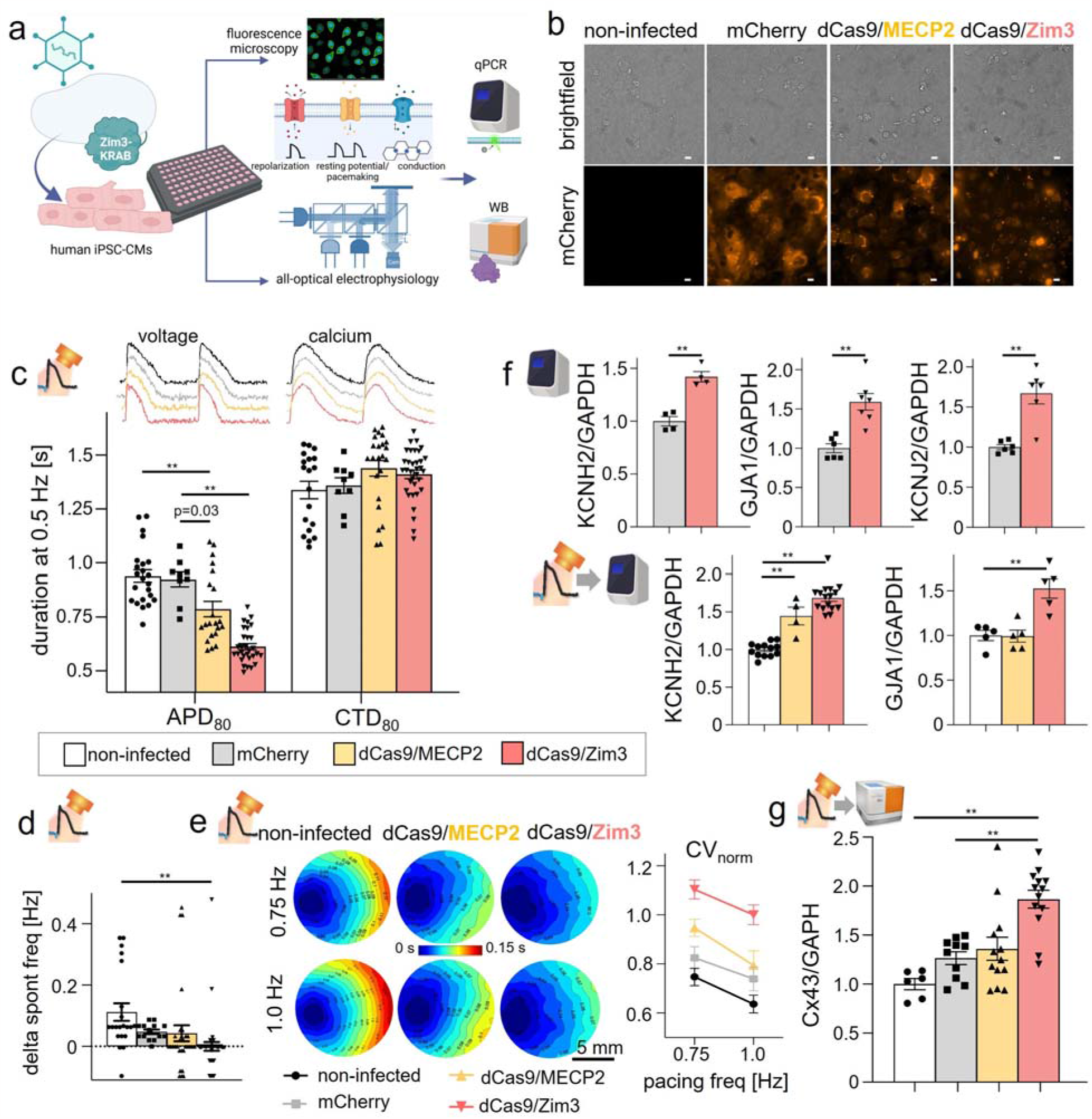
Effects of dCas9/Zim3 expression in human iPSC-CMs: RNA, protein, and electrophysiological consequences. a. Experimental workflow with all-optical electrophysiology(9) and molecular characterization. b. Expression of mCherry tagged constructs (mCherry, MeCP2-KRAB-dCa9-mCherry, Zim3-KRAB-dCa9-mCherry) in human iPSC-CMs. Scale bar 10 µm. c. Functional results from all-optical electrophysiology: changes in APD and CTD under 0.5 Hz pacing (n=9-34 biologically independent samples). d. Relative changes in spontaneous frequency (delta) with respect to the mean for the dCas9/Zim3 group for each run (n=14-40 biologically independent samples per group). e. Changes in conduction shown in example activation maps and as normalized conduction velocity (CV_norm_) at 0.75 Hz and 1.0 Hz pacing (n=2-9 biologically independent samples). Data shown as mean ± S.E.M. f. Molecular analysis (qPCR) in pristine samples (top) and after all-optical electrophysiology (bottom) - KCNH2, GJA1, and KCHJ2 (n=4-16 biologically independent samples, n=3 technical replicates). g. Western blot (Wes by ProteinSimple) of Cx43 post all-optical electrophysiology (n=6-13 biologically independent samples). Statistical comparisons were done using one-way ANOVA with post-hoc Tukey correction. In all cases **p<0.01. Biorender was used.

These gene expression changes and functional effects cannot be attributed to the expression of dCas9, as we have shown that regular KRAB-dCas9 (using the KRAB domain of KOX1/ZNF10) without gRNAs does not alter these genes(6). The fluorescent reporter, mCherry, also does not have such effects. Included here earlier-generation effector construct for CRISPRi – MeCP2-KRAB-dCas9 (a non-KRAB effector, MeCP2, combined with KOX1/ZNF10)(7) caused similar, but much milder changes when delivered in the same type of viral vector - adenovirus with an SFFV (spleen focus-forming virus) promoter and an mCherry reporter, at the same multiplicity of infection, MOI 1000, as Zim3-KRAB-dCas9. Interestingly, MeCP2 has been shown previously to act not only as a transcriptional repressor but also as an activator for many genes(8). No such studies have been conducted with Zim3/ZNF657/ZNF264. It is possible that some but not all KRAB domains or transcription effectors, alone, or when linked to dCas9 can exert the effects seen here. We cannot exclude the SFFV promoter as a possible culprit – it has rarely been used with cardiomyocytes before and is shared by these two constructs (not used in the original KRAB-dCas9 delivery in (6)). Overall, we are unaware of any single perturbation acting as a potent agonist for all three genes. These findings should be taken into consideration when performing CRISPRi studies in iPSC-CMs. Zim3 also needs to be investigated further as a potential therapeutic tool for iPSC-CMs maturation for their use in drug screening, disease modeling and regenerative applications; furthermore, it would be interesting to explore if Zim3 may exert similar effects *in vivo* in the adult human heart as a therapeutic tool.

## Author contributions

JLH and EE designed the experiments; YWH built some of the imaging tools needed; JLH and YWH performed most experiments and analyzed the data, MRP and WL conducted and analyzed some of the functional experiments. EE and JLH wrote the paper.

## Data availability

All data are included in the presented figure.

